# Vectorial Channeling as a Mechanism for Translational Control by Functional Prions and Condensates

**DOI:** 10.1101/2021.08.19.457025

**Authors:** Xinyu Gu, Nicholas P. Schafer, Peter G. Wolynes

## Abstract

Translation of messenger RNA is regulated through a diverse set of RNA-binding proteins. A significant fraction of RNA-binding proteins contain prion-like domains which form functional prions. This raises the question of how prions can play a role in translational control. Local control of translation in dendritic spines by prions has been invoked in the mechanism of synaptic plasticity and memory. We show how channeling through diffusion and processive translation cooperate in highly ordered mRNA/prion aggregates as well as in less ordered mRNA/protein condensates depending on their sub-structure. We show the direction of translational control, whether it is repressive or activating, depends on the polarity of the mRNA distribution in mRNA/prion assemblies which determines whether vectorial channeling can enhance recycling of ribosomes. Our model also addresses the effect of changes of substrate concentration in assemblies that have been suggested previously to explain translation control by assemblies through the introduction of a potential of mean force biasing diffusion of ribosomes inside the assemblies. The results from the model are compared with the experimental data on translational control by two functional RNA-binding prions, CPEB involved in memory and Rim4 involved in gametogenesis.

**Significance Statement:** mRNA/protein assemblies such as functional prions and condensates are involved in locally regulating translation in eukaryotic cells. The mode of regulation depends on the structure of these assemblies. We show that the vectorial processive nature of translation can couple to transport via diffusion so as to repress or activate translation depending on the structure of the RNA protein assembly. We find that multiple factors including diffusivity changes and free energy biases in the assemblies can regulate the translation rate of mRNA by changing the balance between substrate recycling and competition between mRNAs. We mainly focus on the example of CPEB, a functional prion that has been implicated in the mechanism of synaptic plasticity of neurons and in memory.

---

Despite the acknowledged beauty and complexity of the cell’s structure, picturing the cell as a “bag of enzymes” remains a seductively powerful paradigm in biochemistry because most of the information content of the cell resides near the atomic-scale. Much of the major work of the cell, however, takes place on length scales intermediate between the atom and the organelle, a size range that is only now becoming accessible to direct visualization (1). In recent years, cell biologists have begun to suggest that many aspects of biological regulation are also under control at these intermediate length scales both in the nucleus and in the cytoplasm (2). Liquid-like RNA protein condensates form in cells and seem to be involved in gene regulation within the nucleus and also seem to control the translation of RNA sequences into protein in the cytoplasm. Several more ordered aggregates based on amyloid-like assembly, so called “functional prions”, feature in the local control of messenger RNA translation and have been invoked as key players in the molecular basis of memory (3, 4). In this paper, we will explore how the local structures of such aggregates can modulate information processing steps, such as translation, by coupling the vectorial processive dynamics of the ribosomes with their transport through diffusion in the structured environment provided by aggregates. Coupling distinct biochemical reactions through diffusive transport has been called “channeling”(5, 6). Channeling was originally suggested in the context of enzymatic biochemistry as a route by which several enzymes can cooperate to carry out a series of processes efficiently, without losing any possibly unstable intermediate products to the larger cellular environment. Quantitative studies show that, in the enzymatic context, the effects of channeling are significant but modest in magnitude (7–10). Here we will show that the effects of channeling can control translation and that, due to the vectorial processivity of translation, the nature of the control (repression or activation) depends on the details of the aggregate structure. In this way, the localized formation of structured aggregates can control where in the cell RNA messages are translated into protein.

Prion domain detection algorithms have uncovered 240 prion-coding genes (11) out of the 20,000 protein-coding genes in the human genome (1.2%). Seventy-two (11) of these 240 proteins (30%) are also identified as RNA-binding proteins (RBPs). 3.5% of the human proteome contains RNA-binding/processing domains(12). This significant overlap between proteins that can form prions and RNA-binding proteins suggests that prion formation may be rather generally involved in translational control. Some of these prion-like ribonucleic acid binding proteins form well defined amyloid-like assemblies(13). Other ribonucleic acid binding proteins assemble in an apparently more disordered fashion into what are called condensates or “membraneless organelles” like processing bodies (P bodies) (14).

The cytoplasmic polyadenylation element binding protein (CPEB) family provides a notable specific example of a functional prion. Several homologs in the CPEB family, such as CPEB3 in vertebrates(3), Orb2 in Drosophila(16), and ApCPEB in Aplysia(17, 18) can form stable amyloid aggregates in dendritic spines, where they are involved in the maintenance of long-term memory in neurons. In the spines these proteins can form prions when assembled with actin and they also control translation of RNA into synaptic proteins and cytoskeletal proteins, such as actin, forming a feedback loop between the cytoskeleton growth and protein synthesis(3, 4). When CPEB undergoes a transition from its monomeric form to an amyloid aggregate, its regulatory function switches from one of translational repression to one of activation(19). Exactly how this happens remains mysterious. One view is that monomeric CPEB represses translation by recruiting target mRNAs to P bodies, while then forming the more ordered prion simply releases this repression by segregation (20). It has also been proposed that some heterologous translational repressors or activators may be involved in the change of regulation by either recognizing the monomeric or the amyloid form of CPEB (15, 21). Recently, the Cryo-EM structure of Orb2 aggregates has been solved and the geometry of the mRNA/prion aggregate has been uncovered (15). Intrigued by this known aggregate structure, here we suggest that the vectorial nature of mRNA processing, along with the structurally polarized nature of the mRNA/prion aggregates, potentially contributes to the translational activity of CPEB aggregates. Essentially, the prion forms a local translation factory assembly line in which ribosomes are more efficiently recycled. To understand how this might work, recall that the CPEB RNA-binding domain binds to the 3’ end of the target mRNAs(19). This is where translation is terminated. As illustrated in Fig. 1A, during translation ribosomes translocate along an mRNA, starting from the 5’ end, which is found in the outer layer of the mRNA/prion aggregate, moving inwards towards the 3’ end, which is found at the center of prion fiber. After the termination of translation, free ribosomes are released from the mRNA 3’ end and now have the possibility of subsequently being efficiently recruited by a 5’ end of the same mRNA or an alternate mRNA again. Only after the ribosomes have been reused several times will they diffuse away from the prion factory. In fact, envisioning a similar mechanism, it was proposed two decades ago that the circularization of mRNA also could result in a ribosome recycling effect by employing several times a single circular mRNA (22, 23). Ribosome recycling will enhance the translation rate and resembles the substrate channeling effect suggested to occur in various metabolic pathways (24). In translation activation, the ribosomes and the mRNAs play analogous roles as the substrates and the enzymes do in metabolic enzyme assemblies, respectively.

**Fig. 1.**
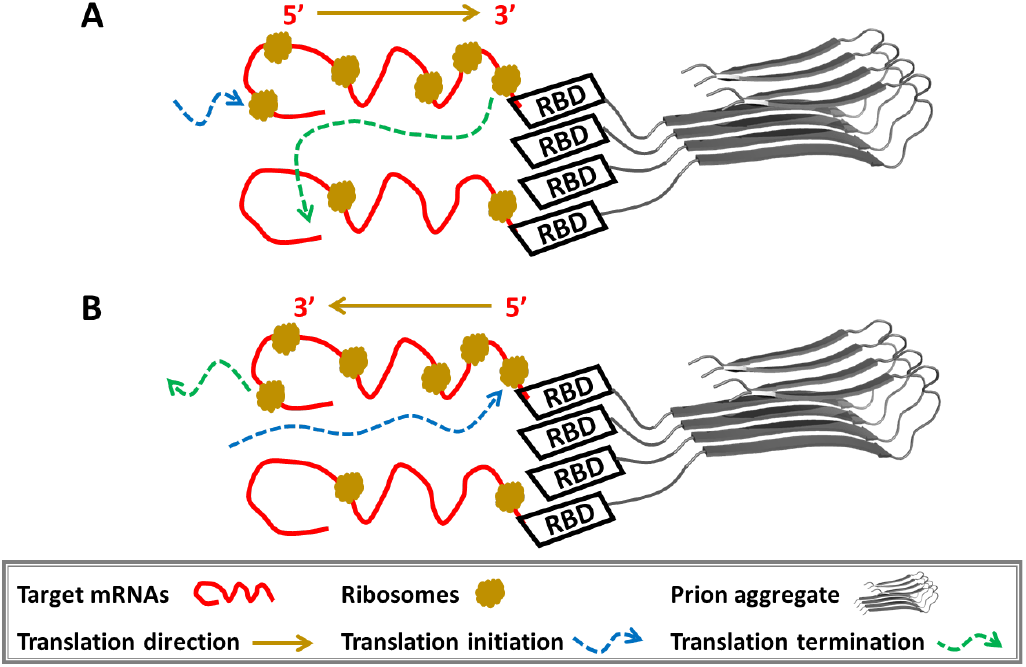
The organized structure of the mRNA/prion aggregates leads to the regulation of translation activity. **A**. Diagram for the assembly of mRNAs and Orb2/CPEB aggregates. The prion core of the Orb2 aggregates adopts a trimer ribbon conformation, located at the center of the Orb2 fiber(15). Here we use only one monomer ribbon to represent the fiber structure. The rest of the Orb2 protein binds to the 3’ UTR of target mRNAs using an RNA-binding domain (RBD). **B**. The structure of the mRNA and Rim4 aggregates displays the opposite polarity. Rim4 RBD binds with the 5’ UTR of target mRNAs. Notations are shown in the lower panel.

Another example of an RNA-binding prion which adopts an amyloid-like structure is provided by the regulator of IME2 kinase, Rim4. Interestingly, the Rim4 amyloid-like aggregate is reported to repress the translation of its targets(25), rather than activate them as in the CPEB case. In contrast with CPEB, the Rim4 protein binds to the 5’ end of its target mRNAs. This initiation site is now buried deep in the assembly, making it less accessible to ribosomes. As shown in Fig. 1B, in this vectorial geometry, upon terminating translation, ribosomes are able to diffuse directly into the bulk aqueous phase of the cell, thereby lowering the probability that they can be efficiently reused, quite the opposite of what we see for the CPEB case. What’s more, we would expect the translation initiation to slow down since the ribosomes have to diffuse through the dense mRNA layer of the assembly to find their target start codon at the buried 5’ end.

We propose that the vectorial nature of translation, coupled with the polarized structure of RNA-binding prion aggregates determines whether an assembly represses or conversely activates translation. In this study, to explore this idea we have modelled the translation process in RNA/prion systems by coupling a simplified description of the totally asymmetric simple exclusion process (TASEP) model for the transport of bound ribosomes on mRNAs (26, 27) with the reaction-diffusion equation for free ribosomes. In steady state, the coupled equations turn into a Poisson equation for diffusion with many pairs of sinks and sources. For aggregates with spherical geometries, the Green’s functions for this Poisson equation can be found analytically using the well known electrostatic analogy (28). But for more complicated geometries like a cylindrical structure, which is closer to the real prion fiber structure, the Green’s functions may also be found numerically using finite element analysis(29). We explore these steady state solutions for various geometries of the assemblies, and spatial diffusivity patterns within them. Assemblies may also attract by binding or sterically repel ribosomes. Therefore we have also studied the effects of a free energy bias for ribosomes to remain in the assembly or to be pushed out. The free energy bias will reflect the balance between excluded volume and specific ribosome binding to the assembly. The relative translation rates are then calculated to quantify how translational control is determined by the size of aggregates, the inhomogeneous nature of the diffusivity through the assemblies, and the free energy bias for free ribosomes to be found inside versus outside the assemblies. For CPEB and Rim4 aggregates, the direction of translational control (activation or repression) of the channeling effect is correctly predicted by the present vectorial channeling model. Of course the vectorial channeling hypothesis, while perhapsbeing a common mechanism, doesn’t exclude, but rather may cooperate with alternative mechanisms of translational control. The consistency between the structures of assemblies and their functions in translation regulation suggests that RNA-binding prions and UTR regions of their target mRNAs have coevolved to form large-scale structures that can function as organized translation factories.

We have also applied our algorithm to less ordered assemblies to provide a zeroth order description of the translational control effects of nonamyloidogenic RNA/prion condensates. In contrast with the amyloid-like assemblies, fully non-polar condensates show no vectorial “channeling” effect on the average in the absence of a free energy bias raising ribosome concentration in the condensate. We show, however, that specific internal structures of the condensates would provide local polarity to regulate the translation of mRNAs inside the condensates. This raises the issue of the degree of structural order in condensates. The present model suggests that the direction and significance of translational control of assorted RNA/protein assemblies can be cooperatively regulated by the vectorial channeling effect through modifying assembly structure, along with the effect of substrate concentration changes in assemblies and the effects of substrate competition within them.

## Model

Vectorial channeling involves the interplay of the processive kinetics of translation and diffusion in a structured three dimensional space. The peptide synthesis rate from the processive translation is determined by the number of bound ribosomes and the net translocation rate of ribosomes along the mRNAs. Previous studies on translation in Yeast cells(26) suggest that the rate-limiting step in translation is initiation and that ribosome collisions are rare. In the present model we therefore make a practical assumption that the translocation rate of bound ribosomes along the mRNAs is a fixed constant and thus that the translation rate is proportional to the number of bound ribosomes per mRNA. With this assumption, the TASEP model simplifies to give the rate of change of bound ribosome number on the i*th* mRNA:

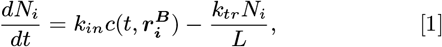

where *L* is the codon length of the i*th* mRNA and *N_i_* is the number of bound ribosomes on i*th* mRNA. The lengths of the target mRNAs will be assumed uniform. As shown in Fig. 2A, we see the increase of bound ribosome number is proportional to the concentration of the free ribosomes at the start codon, 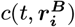, with an initiation coefficient *k_in_*. The decrease of bound ribosome number at termination is just the product of the probability of finding a ribosome at the stop codon, *N_i_/L*, and the ribosome translocation rate, *k_tr_* (26) when there are no traffic jams.

**Fig. 2.**
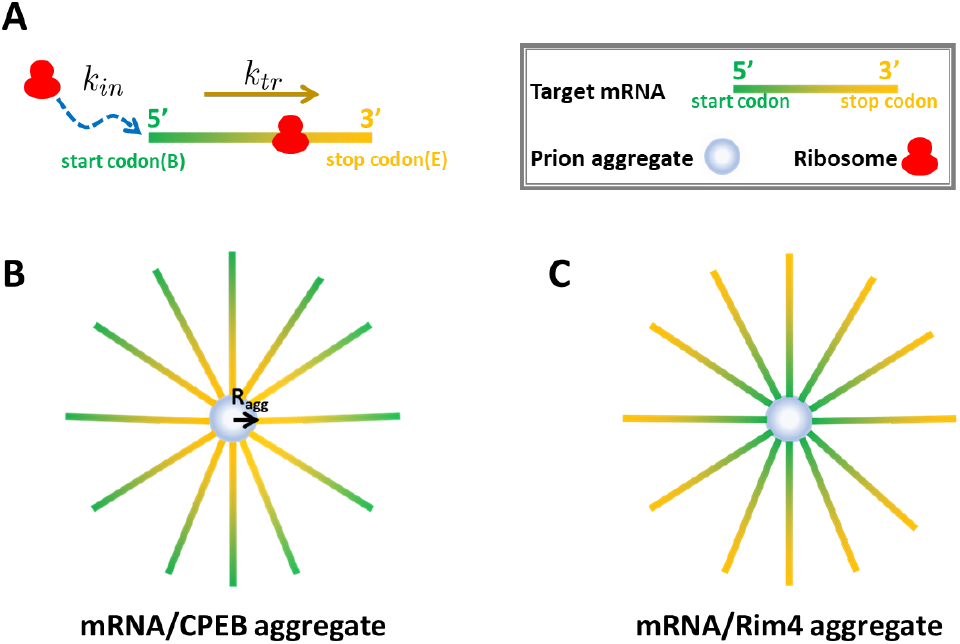
**A**. Kinetics of translation on a single mRNA in the simplified TASEP model depends on the constant translation initiation rate, *k_in_*, and the elongation rate, *k_tr_*. **B**. A sketch of a spherical mRNA/CPEB aggregate. **C**. A sketch of a spherical mRNA/Rim4 aggregate. The prion aggregates are shown as grey inner spheres with a radius of *R_agg_* and each mRNA is represented by a straight line with gradient color from green to yellow, from 5’ end (translation starting end) to 3’ end (traslation stopping end), as noted in the upper-right corner.

The change of the free ribosome concentration, *c*(*t, r*), is governed by a reaction-diffusion equation,

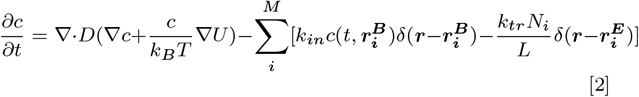

In this equation, *D*(***r***) and *U*(***r***) are the inhomogeneous spatially varying diffusion coefficient and the potential of mean force for free ribosomes to move in the vicinity of the assembly. *M* is the total number of mRNAs bound in the aggregate. The start codons of the mRNAs act as sinks for ribosomes while the stop codons act as sources. We consider codons to be sufficiently small so as to act as point sinks or sources at 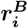 or 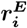, The source *N_i_* is obtained from Eq. 1. We assume an asymptotic boundary condition for the ribosome pool, *c*|_*r*→∞_ = *c*_0_, where *c*_0_ is a fixed constant.

### Stationary solution

In steady state, *N* and *c* are independent of time, so the time derivatives vanish. Eq. 1 then gives:

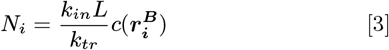

We see that the number of bound ribosomes is proportional to the free ribosome concentration at start codons, 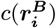. Substituting this result into Eq. 2, we obtain a coupled reaction diffusion equation:

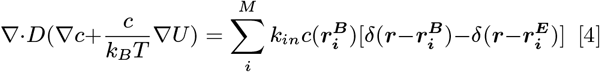

If there were no sources or sinks, the solution of this equation is *c*(***r***) = *c*_0_*e*^-*U/k_B_T*^. The additional delta function sources and sinks then give an implicit equation

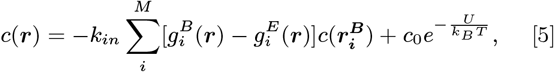

where 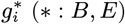 is the Green’s function of the steady state Smoluchowski equation:

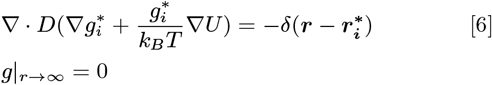

The effective translation rate of the j*th* mRNA depends on 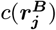, which follows from Eq. 5. The self-term, 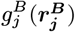, in that equation reflects a renormalization of the initiation rate by diffusion, which has already been included in *k_in_*. Dropping this self-term in Eq. 5 thus gives

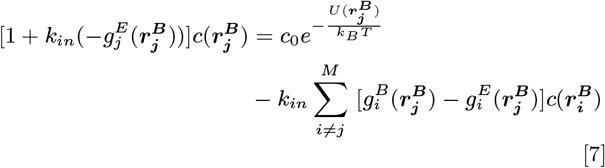

This equation can be organized into a matrix form:

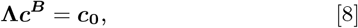

where **Λ** is a coefficient matrix with elements:

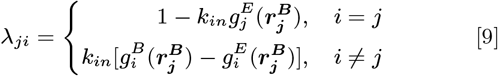

***c^B^*** and ***c***_**0**_ are two vectors whose i*th* elements are 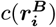 and 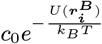 respectively for i ranging from 1 to M, the total number of mRNAs. One then finds the vector of effective initiation concentrations:

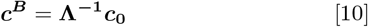

For the perfectly symmetric sphere model sketched in Fig. 2B/C, each mRNA is identically situated thus 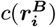 is the same for all i, denoted as *c*(***r^B^***). In this case, we find explicitly:

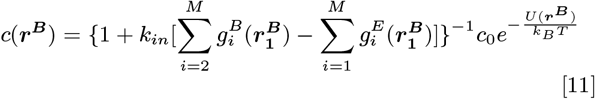

In the absence of the assembly, there will be no potential of mean force bias (*U* = 0) and no inhomogeneity in diffusivity. In that case the concentration of free ribosomes at the start codon of the single mRNA, 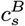, is given by

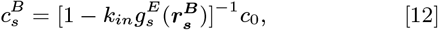

in which the full Green’s function 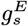 enters. Using Eq. 12 and Eq. 10 we can evaluate the translation regulation effect as a ratio, 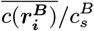 (or 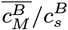 for short) to determine the direction (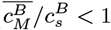 for repression; 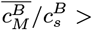 for activation) and the extent of the translational control.

#### Model potential of mean force

For simplicity we will use a smooth step function as a potential of mean force, *U*(*r*), to study the effect of any free energy difference for ribosomes being inside of the assembly versus in the environment.

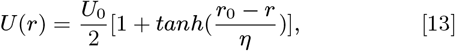

where *r* is the distance from the center of the assembly. *U*_0_, *r*_0_ and *η* are parameters regulating the strength and shape of *U*.

#### Calculating Green’s functions via electrostatic analogy

When *U* = 0, the steady state Smoluchowski equation Eq. 6 has a simple electrostatic analogy. Its solution reflects the electrostatic potential from a point charge in a dielectric medium. With homogeneous diffusivity, the equivalent dielectric is homogeneous, so we find the Coulomb potential:

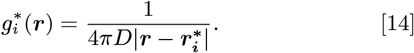

The RNA/prion assemblies, being dense membraneless compartments, will provide an inhomogeneity in the diffusion coefficient for free ribosomes. For the full spherical inhomogeneous geometry, when *U* = 0, one can still find a straightforward electrostatic analogy for the Green’s function - a point charge inside a dielectric sphere. Using the method of images in the electrostatic analogy, the solution of Eq. 6 inside the assemblies for free diffusion without a potential of mean force is:

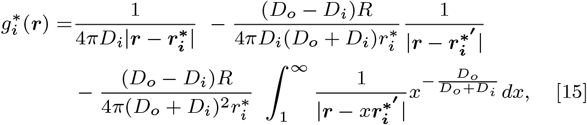

where *D_i_* and *D_o_* are the diffusion coefficients inside and outside the spherical assembly and *R* is the radius of the assembly. 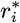 is the length of vector 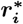 and 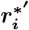 is the mirror image of 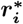 satisfying:

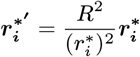

For the cylindrical assemblies and for systems with a non-zero potential of mean force, *U* ≠ 0, we used COMSOL Multiphysics(29) to find the corresponding Green’s functions numerically.

## Results

We illustrate our results for several different structural geometries and situations. For this purpose, we have used several sets of values of the parameters which we describe first. The values of these basic parameters have been organized in Table.1. In the present model, mRNAs are considered to be sticks connecting the start codons and stop codons. The length of the mRNA sticks, *l*, is considered uniform for all the mRNAs in one system. For the homogeneous spherical aggregate model, we have varied *l* over a realistic range from 50 nm to 130 nm (as shown in Fig. S2). Shortening the distance between the start codon and the stop codon facilitates the recycling of ribosomes and increases the translation rate slightly. We fixed *l* at 50 nm for further comparisons. For the cylindrical aggregates, another structural parameter, the linear density of mRNAs is also needed. We have studied several values of the linear density for the homogeneous cylindrical model (as shown in Fig. S3) and for most of the comparisons we have selected a linear density of 1 mRNA/nm for the other cylindrical models. For the radius of the spherical or the cylindrical prion core of the assemblies, we estimate a value of 10 nm based on the EM picture of the Orb2 fiber(15). Although the elongation rate of bound ribosomes can be measured by ribosome profiling experiments, the translation initiation rate is more challenging to define and measure experimentally. We will assume the start codon provides spherical ribosome sink with a small radius of *ϵ*. Then the diffusion limit of the kinetic rate of translation initiation, *k_D_*, equals 4*πD_o_ϵ*. If the initiation rate were larger, *k_in_* ≥ *k_D_*, the translation initiation process would become diffusion-controlled(28). The rate of translation initiation may be smaller and thus not be diffusion-controlled. The length of a start codon, 2*ϵ*, is about 1 nm(33), so *k_D_* is around 2 × 10^6^ *nm*^3^/*s*.

**Table 1.** Model parameters.

| Parameter | Value |
| --- | --- |
| $R_{agg}$, radius of prion aggregate core(15) | 10 nm |
| $k_{tr}$, ribosome translocation rate on mRNA(26, 30) | 2.63 ~ 6.8 aa/s |
| $D_o$, diffusion coefficient of free ribosomes in cytoplasm (31) | 0.3 $\mu m^2/s$ |
| Range of ribosomes level, $c_0$, among different tissues (32) | 0.6 ~ 4.7 $\mu M$ |

aa: amino acid; <sup>[a]</sup> We assume all target mRNAs code 500 aa.

### Vectorial channeling is governed by the polarized structures of mRNA/prion aggregates and is amplified by inhomogeneous diffusivity

We will first study situations having diffusion effects alone and set the potential of mean force to zero, *U* = 0. We will then vary the aggregate size of RNA/prion assemblies, *M* and the diffusion coefficients inside assemblies *D_i_* to investigate the effects of structure and inhomogeneous diffusivity on translational control by RNA/prion aggregates. For the symmetric spherical model shown in Fig. 2B/C, we used Eq. 15 to calculate the Green’s functions analytically and then obtained the translation rate using Eq. 11. For the inhomogeneous cylindrical model, we used COMSOL Multiphysics to calculate numerically an approximation to the Green’s functions for the point sources inside an infinitely long cylinder and then we used the resulting Green’s functions as approximations in the finite cylinder models to solve Eq. 8 for the net translation rate. As shown in Fig. 3, the polarized spherical assemblies are more compact and are more efficient in vectorial channeling than the cylindrical assemblies. Fig. 3A/B shows the results when initiation is assumed to be diffusion limited, while Fig. 3C/D shows the results for slower initiation. The results show that translation is predicted to be enhanced for the CPEB case but that translation is repressed for the Rim4 geometry. As we mentioned in the introduction, the translational enhancement for CPEB arises from efficient ribosome recycling while the translational repression for Rim4 arises from the difficulty of free ribosomes to find a path towards the start codons, which are all buried together at the center of the assemblies. As the aggregates grow in size, the structural effects become more prominent and amplify the translational control. A smaller diffusion coefficient inside the assemblies, *D_i_*, leads to greater translational control. This results from the impact of the spatially inhomogeneous diffusion coefficient mentioned above: substituting Eq. 14 into Eq. 12 shows the translation rate of one single mRNA is proportional to [1 – *k_in_/*(4*πDl*)]^-1^. It is easy to see that lowering the diffusion coefficient increases ribosome recycling from the stop codon and therefore accelerates translation; Similarly, if there are two sinks separated by *l* in distance, the substrate concentration at one sink is then proportional to [1 + *k_in_*/(4*πDl*)]^-1^, suggesting that lowering the diffusion coefficient would increase the competition for ribosomes between start codons and would slow down translation. Likewise increasing *k_in_* exaggerates translational control by aggregate structure. As shown in Fig. 3, the translational control by the assemblies is more significant when *k_in_* = 2 × 10^6^ *nm*^3^/*s* (the diffusion limit) than when *k_in_* = 2 × 10^5^ *nm*^3^/*s* (slow initiation). In the latter case, the translation is mainly limited by the small initiation rate so the other factors only cause subtle changes. As shown in Fig. 3A, the limit of the activation effect in the CPEB model is an increase 1.3 fold when *k_in_* = 2 × 10^6^ *nm*^3^/*s*. The activation effect could be increased by up to 3.4 fold when *k_in_* = 6 × 10^6^ *nm*^3^/*s* (as shown in Fig. S4). For Rim4, the limit of the repression effect is 2 × 10^-3^ fold when *k_in_* = 2 × 10^6^ *nm*^3^/*s*, as shown in Fig. 3B. The repression effect would be increased by up to 7 × 10^-4^ fold when *k_in_* =6 × 10^6^ *nm*^3^/*s*.

**Fig. 3.**
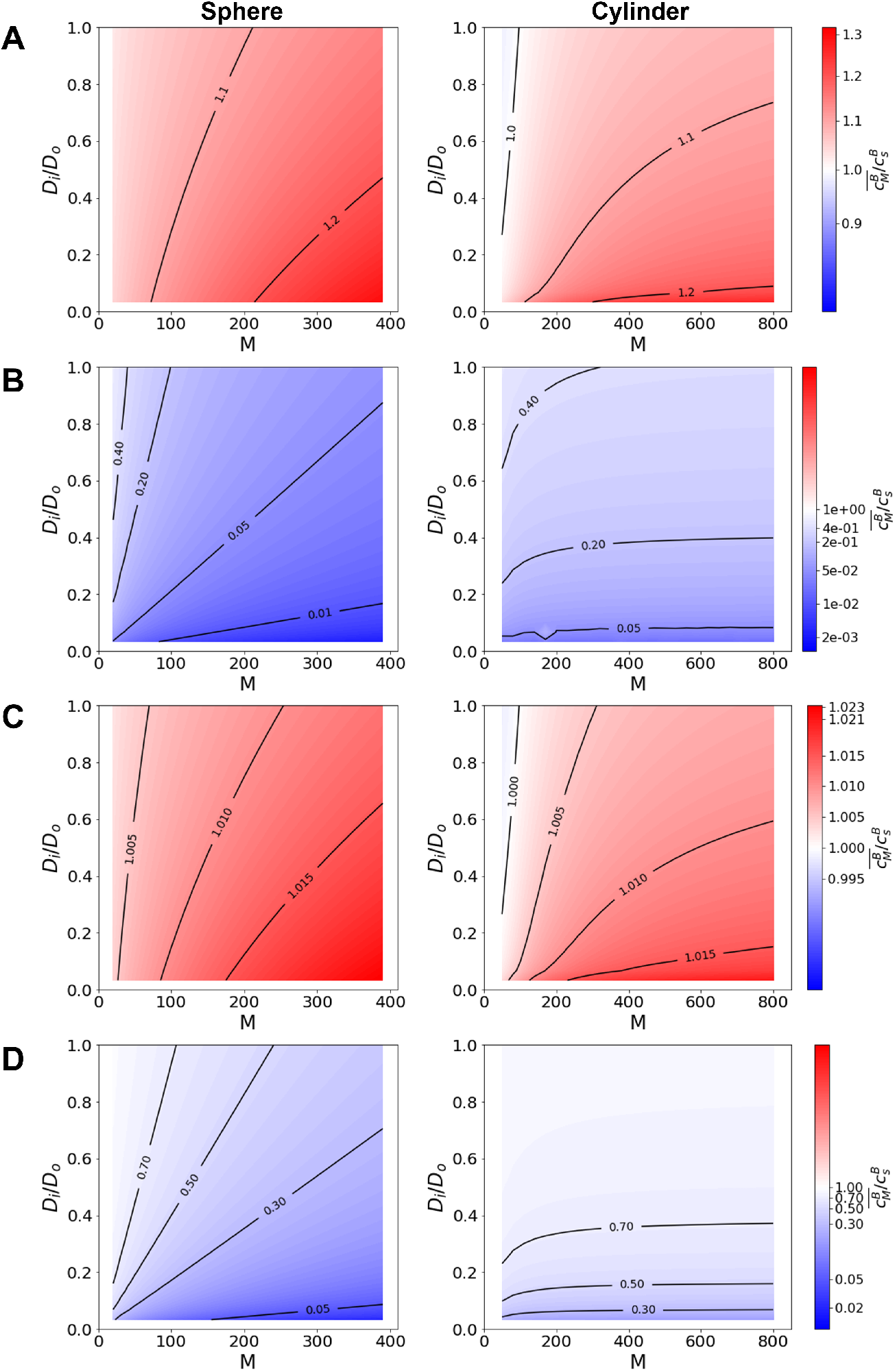
The structure and the inhomogeneity of the diffusion coefficient within the mRNA/prion assembly determines the direction and extent of translational control. The colors in these 2D maps indicate the relative translation rate 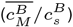, ranging from blue to red that is from translational repression to activation. The x axis, *M*, is the aggregate size and the y axis, *D_i_*/*D_o_* is the ratio of the diffusion coefficient inside the assembly versus outside the assembly. **Left panel**: the spherical aggregate model. **Right panel**: the cylindrical aggregate model with an mRNA linear density of 1 *nm*^-1^. **A** or **B**: Results for the CPEB or Rim4 model under diffusion-controlled condition (*k_in_* = 2 × 10^6^ *nm*^3^/*s*). **C** or **D**: Results for the CPEB or Rim4 model when initiation is slower than the diffusion limit (*k_in_* = 2 × 10^5^ *nm*^3^/*s*).

### The potential of mean force inside mRNA/prion aggregate regulates translation via modifying substrate concentration

As well as providing spatial heterogeneity in diffusion, the RNA/prion assemblies may provide a free energy barrier at the boundary of the assemblies for entry or may attract ribosomes to the assembly interior. We focus on spherical assemblies with spherical potentials of mean force *U*(*r*). The precise Green’s functions for this case were computed numerically in COMSOL Multiphysics. The potential *U*, given in Eq. 13, is close to 0 far away from the assemblies and is close to *U*_0_ at the center of the assemblies. The parameters *r*_0_ and *η*, which determine the exact location and span of the barrier, were selected to make the potential barrier shell start from the outer boundary of the assemblies and to have a width of half of radius of the assemblies (as shown in Fig. S1A). We varied the aggregate size, *M* and the free energy *U*_0_ to investigate the effect of binding free energy difference on the translational control by RNA/prion aggregates. Both the diffusion limited (Fig. 4A/B) and slow initiation (Fig. 4C/D) situations were studied and both the homogeneous and the heterogeneous diffusivity were tested. Not surprisingly, our results indicate that an attractive potential for free ribosomes that guides them into the assemblies (*U*_0_ < 0) increases the translation rate by concentrating ribosomes within the assemblies while a repulsive potential (*U*_0_ > 0) slows down translation. For our specific *U* function, the potential at the start codons is around 0 for the CPEB case, but around *U*_0_ for the Rim4 case. So, in the CPEB case, the substrate concentrating effect caused by the potential is almost negligible compared with the vectorial channeling effect, as shown in Fig. 4A. These two effects are more comparable when initiation is slow and the channeling effect is thus also subtle, as shown in Fig. 4C. In contrast, the free energy differences yield a more dramatic effect for the Rim4 case as shown in Fig. 4B/D. It should be noted that we have only taken one specific potential of mean force for illustration. Modifications to the parameters and the detailed form of *U*(*r*) will vary the results quantitatively. For example, if the location of the barrier is moved outwards, the potential at mRNA start codons in CPEB case would be closer to *U*_0_ instead of 0 and the substrate concentration effect would be much more significant.

**Fig. 4.**
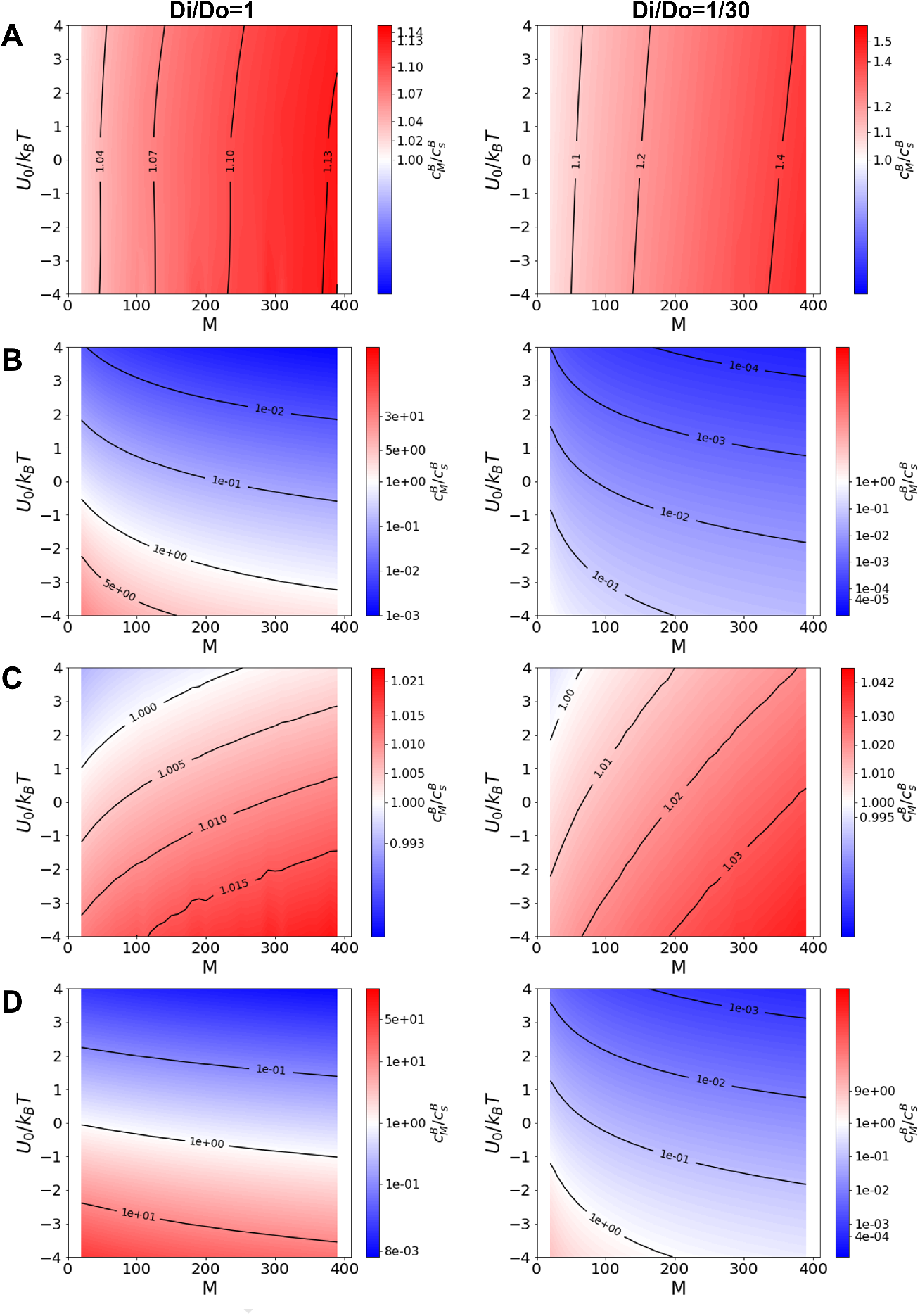
An attractive potential of mean force for ribosomes inside the mRNA/prion assembly increases the translation rate while a repulsive potential lowers the rate. Here we only studied the spherical model. The setups for the color codes and the x axis are the same as for Figure. 3, while the y axis, *U_o_*/*k_B_T* represents the free energy barrier raised by the assembly. A negative value of *U_o_*/*k_B_T* means there is an attractive potential inside the assembly, while a positive value means a repulsive potential. **Left panel**: Results with a homogeneous diffusivity (*D_i_*/*D_o_* = 1). **Right panel**: Results with an inhomogeneous diffusivity (*D_i_*/*D_o_* = 1/30). **A** or **B**: Results for the CPEB or Rim4 model under diffusion-controlled condition (*k_in_* = 2 × 10^6^ *nm*^3^/*s*). **C** or **D**: Results for the CPEB or Rim4 model when initiation is slower than the diffusion limit (*k_in_* =2 × 10^5^ *nm*^3^/*s*).

### The internal architecture of condensates regulates their translational control

The vectorial nature of channeling relies on the locally polarized structure of the aggregates. For functional prions the polarized nature of the structure is clear. To investigate the translational control by disordered nonamyloidogenic RNA/prion condensates, we must understand more about their internal structure. To explore this, we first generated randomly distributed and oriented mRNAs in a large sphere whose radius is 500 nm. We investigated only the pure diffusion limited case where we expect the most significant effect. We also neglect the effect of the potential of mean force with in the condensate to focus on the vectorial aspects. Eq. 15 provides the Green’s functions analytically and then the matrix Eq. 8 is solved to obtain effective translation rate for each mRNA. Unlike the amyloid RNA/prion assemblies which have clearly polarized mRNA distributions, for the non-polarized condensates the effective translation rate is nearly independent of the mRNA copy number in the condensates. When the diffusivity is homogeneous (*D_o_*/*D_i_* = 1), the non-polarized condensates provide no translational control since for each mRNA, the influences from other mRNAs cancel with each other, as the solid black line shows in Fig. 5. When the diffusion coefficient is small in such a large condensate, however, the ribosome recycling from a single mRNA to itself would be more significant and therefore enhance the translation, as the solid lines show in Fig. 5.

**Fig. 5.**
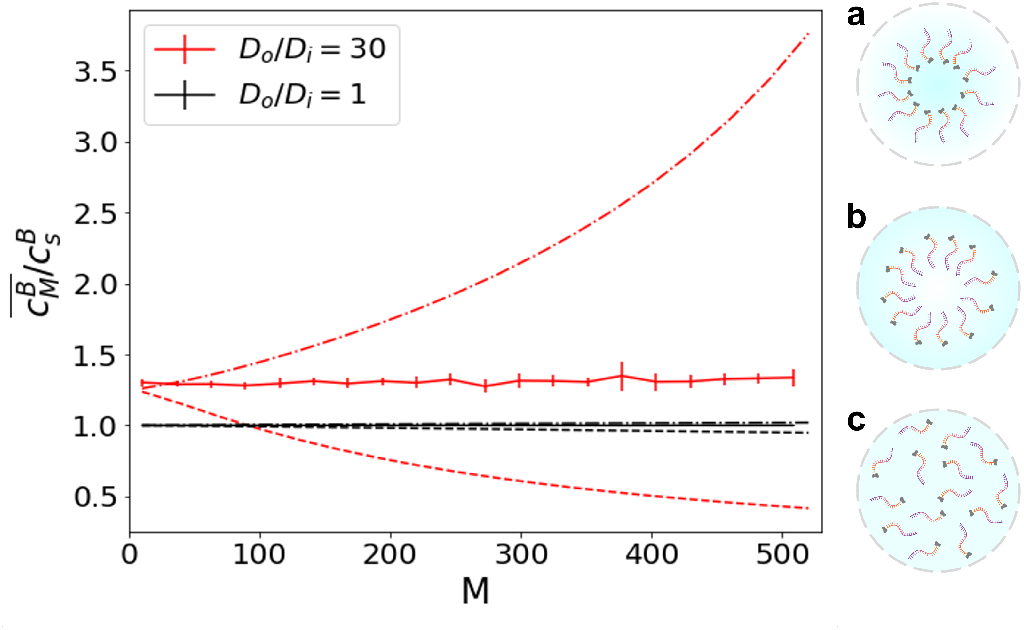
Substructures of condensate determine the polarity of mRNA/prion assembly and therefore regulate its translational control. Here we take condensates containing CPEB and its target mRNAs as an example. The average translation rate, 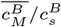, for spherical condensates with a radius of 500 nm is plotted against mRNA number in the condensates, *M*, under diffusion limited condition (*k_in_* = 2 × 10^6^ *nm*^3^/*s*). Several internal architectures of condensates are tested when *D_o_/D_i_* equals to 1 (shown as black lines) or 30 (shown as red lines). Monomeric CPEB (colored in grey) binds with the 3’ end of its target mRNA (colored from purple to orange, from 5’ to 3’). **a**: condensate with an RNA shell with an outer radius of 450 nm. (shown as dot-dashed lines) **b**: condensate with an RNA core with an outer radius of 450 nm. (shown as dashed lines) **c**: completely disordered condensate. (shown as solid lines) 20 random mRNA distributions are generated for each data points in case **c** to obtain the mean values and error bars.

Condensates with diverse internal architectures have been observed that support various biological functions(34). Condensate scaffolds, composed of RNA molecules and various nucleoproteins may behave like the porous polymers(2). Ribosomes, being large ribonucleoprotein complexes, interact with the RNA scaffold through steric exclusion and possible binding and can be trapped in such porous structures. The low activities of large ribosome molecules in RNA scaffold may result in a repulsive potential of mean force for ribosomes inside some condensates. This would result in a translation repression effect based on the analysis in the last section. At the same time, specific substructures of condensates might possess local polarity and thus regulate translation by vectorial channeling. For condensates containing monomeric CPEB binding with its target mRNAs, an architecture with an RNA shell and protein core has a similar polarity as the CPEB prion aggregate which then would enhance the translation of CPEB’s target mRNA as shown in Fig. 5a. In contrast, a condensate architecture with an RNA core and protein shell gives a reversed polarity and therefore represses the translation of CPEB’s target mRNA as shown in Fig. 5b. For proteins binding with the 5’ end of their target mRNAs like Rim4, the results for these two cases would be simply reversed. We have also investigated a model for the condensate containing many distinct RNA/prion oligomers and noticed that the direction of translational control is largely determined by the polarity of the oligomers themselves in this case as shown in Fig. S7.

High oligomer density in the condensate increases substrate competition and thus slightly slows down translation rate as shown in Fig. S8.

## Discussion

### The limitations of the present model

The “traffic jam” effect that can result from ribosome crowding on mRNAs which would curb elongation is not considered in the present model. Though experimental measurements on protein synthesis rate and ribosome footprinting in Yeast cells(26) did not show any evidence for ribosome crowding, the traffic jam effect might still play a role in translational control in other cells and in organelles. The original TASEP model takes account of the collisions of ribosomes caused by crowding on the mRNA by introducing an exclusion between neighboring ribosomes. Numerous theoretical studies (33, 35, 36) have been developed on this basis. Although the quantitative analysis for the influence of traffic jams in aggregates is beyond the scope of our model, we comment briefly on the regime in the present model where the traffic jam effect can be taken as negligible. Within the reasonable range of parameters, the translation enhancements seen in the present model are less than 4 fold unless there is an extremely strong attractive potential raising concentration in assemblies. Hence, increasing the translation rate in the single mRNA should have a similar magnitude of effect for traffic jams. In the stationary solution, the density of bound ribosomes for single mRNA translation is given by substituting Eq. 12 into Eq. 3: *N/L* = *k_in_ c*_0_ [1 — *k_in_*/(4*πD_o_l*)]^-1^ /*k_tr_*. The value of *k_in_* is around or less than 4*πD_o_ϵ* in our study and *ϵ* ≪ *l*, so *N/L* ~ *k_in_ c*_0_/*k_tr_*. Consistent with the mathematical model established by Lodish(37), the ribosome density on mRNAs, *N/L* would rise up to the crowding limit (~05 rpc, ribosomes per codon)(26) if the ribosome concentration, *c*_0_ and initiation rate, *k_in_* were high enough. From Table. 1, we could select typical values for *c*_0_ of 1*μM*(~ 6 × 10^-7^ *ribosomes/nm*^3^) and *k_tr_* of 5 aa/s. When *k_in_* = 2*e*5 *nm*^3^/*s*, the bound ribosome density is around 0.02 rpc, below the crowding limit. For the high *k_in_* where we are near the diffusion limit, the traffic jam effect may be important but should be modest for specific systems like the dendritic spines that have only a limited pool of available ribosomes(38). In short, when the ribosome abundance is high, the translation of mRNAs with high initiation rate may be lowered because of ribosome crowding on mRNAs, but otherwise the general neglect of traffic jam seems a reasonable assumption.

The ribosome consists of 2 heterogeneous subunits, 60S and 40S for eukaryotes. To initiate translation, the smaller ribosomal subunit (40S) binds to the start codon of mRNA and then recruits the other subunit (60S) to form the ribosome assembly. In the present model, the ribosome is treated as one substrate. It seems that the binding of the small subunit to mRNA is the rate-limiting step, compared with quick recruitment of the large subunit(39). From the perspective of the steady state approximation, the initiation rate of translation will thus be determined by a rate-limiting step, with a rate proportional to the 40S concentration. To take the assembling of ribosomes into account, one only needs to replace the free ribosome concentration into 40S concentration in the model. The kinetics of 60S could also be described by a similar model. It is expected that considering the recruitment of 60S would slightly amplify the translational control.

The production and decay of mRNAs by transcription and ribonucleases were also not taken into account in our calculations, since we assumed that these two factors are balanced with each other in steady state. It would be possible but challenging for our model to address the time dependence and fluctuations of mRNA numbers via solving more complicated time-dependent equations.

## Conclusions

To compare the present model with experimental results, a summary concerning translational control by RNA/prion assemblies is provided in Table.2. As we mentioned before, the natural translation regulation of CPEB is bi-directional and depends upon the structure of mRNA/CPEB assemblies. Our model provides a reasonable explanation for both of the two directions by considering the dominance of either the vectorial channeling effect or the effects of substrate concentration and competition in the assemblies. For the Rim4 case, the model gives proper predictions whether or not initiation is diffusion limited and suggests that the repressive translation regulation of Rim4 mainly arises from the competition of ribosomes between start codons in a dense cluster.

**Table 2.** Translational control by RNA-binding prions aggregates.

| Prion | Aggregate type | RBS<sup>a</sup> | direction<sup>b</sup> | level (expt.)<sup>c</sup> | level (model)<sup>d</sup> |
| --- | --- | --- | --- | --- | --- |
| CPEB | amyloid | 3' UTR | + | ~1.5 folds(19); ~5 folds(15) | 1.3 folds; 3.4 folds |
| Rim4 | amyloid | 5' UTR | - | ~0.1 folds(25) | 0.1 folds |
| CPEB | condensate (P body) | 3' UTR | - | ~0.3 folds(19); ~0.4 folds(20) | 0.4 folds |
| TDP-43 | amyloid | 3' UTR(40) | +(41, 42) | / | / |

<sup>a</sup> mRNA binding site of prion. <sup>b</sup> the direction of translational control. +: activate translation; -: repress translation. <sup>c</sup> the level of translational control measured by experiments. <sup>d</sup> the closest prediction given by the model comparing with the experimental level of translational control.

How general is the vectorial channeling mechanism? We looked for additional examples of functional amyloid-like RNA-binding prions. Unfortunately, because of the difficulties in characterizing prion proteins containing intrinsic disordered sequences, the structures and functions of most RNA-binding prions are still enigmatic. We have only identified a single additional candidate, the TAR DNA-binding protein (TDP-43), which binds with 3’ UTR of its target mRNAs(40). Intriguingly, the translational control by TDP-43 is also bi-directional just as in the CPEB case. It has been proposed that binding with soluble TDP-43 promotes instability and degradation of its target mRNAs and TDP-43 aggregation may impede this inhibition to its targets.(42) While in regenerating muscle, TDP-43 has been reported to form amyloid-like assemblies binding with sarcomeric mRNAs and regulating their translation during skeletal-muscle formation.(41) Further experiments could be designed to test the vectorial channeling hypothesis. A direct test of the vectorial channeling hypothesis can be given by merging the CPEB target sequence to the 5’ UTR or the 3’ UTR of a luciferase mRNA and running a luciferase assay on these two constructs upon CPEB overexpression and aggregation.

In this paper we have contrasted the highly polarized assembly of a prion to more disordered assemblies, which caricature condensates. Our result shows that nontrivial local structure of condensates may give sufficient polarity to make vectorial channeling significant inside condensates. Precise structured modelling of the internal condensate architecture, however, is still challenging. We look forward to a better characterization of the local condensate structures from future experiments and simulations.

We also note that the vectorial channeling hypothesis suggests a natural evolutionary route to higher order structure in the cell. It provides a selection pressure for a “bag of enzymes” to become an exquisitely structured entity.

Consistent with the natural functions of various RNA/prion assemblies, our model suggests that the translational control by RNA/prion assemblies may be regulated by multiple factors including the vectorial channeling effect of polarized structures, substrate (ribosomes) competition, and substrate concentration in assemblies. This model may also be applicable to other spatially vectorial reactions in biological materials, such as the transcription process in chromosomes where compartments are formed. Based on our model, further studies about the polarity of gene distribution, the diffusivity of polymerase in the nucleus, and other chromosome properties at the intermediate length scale would be helpful for investigating transcription processes at the mesoscale.

## ACKNOWLEDGMENTS

This work was supported by the Center for Theoretical Biological Physics, sponsored by NSF grants CHE-1743392 and PHY-2019745. Additionally, we wish to recognize the D.R. Bullard Welch Chair at Rice University, Grant C-0016 (to Wolynes).

