## Supplemental Information for "Vectorial Channeling as a Mechanism for Translational Control by Functional Prions and Condensates"

### Supporting Information Text

#### Using the fibonacci lattice to determine the locations of equally distributed mRNAs

We used a fibonacci lattice to find appropriate coordinates of approximately equally distributed M points on the surface of a sphere with a radius of $R$:

$$\begin{aligned} z_i &= R[1 - \frac{2(i-1)}{M-1}] \\ x_i &= cos(i\pi\phi)\sqrt{R^2 - z_i^2} \\ y_i &= sin(i\pi\phi)\sqrt{R^2 - z_i^2} \end{aligned} \tag{1}$$

or on the side surface of a cylinder with a radius of $R$ and a linear density of $\rho$:

$$\begin{aligned} z_i &= (i-1)\rho^{-1} \\ x_i &= cos(i\pi\phi)R \\ y_i &= sin(i\pi\phi)R \end{aligned} \tag{2}$$

where $(x_i, y_i, z_i)$ is the coordinate for i$th$ point. Here $\phi = 3 - \sqrt{5}$ and i ranges from 1 to M.

#### Comparing the numerical Green's function values with analytical ones

For spherical assemblies without any potential of mean force, Eq. 15 in main text shows the Green's function inside the assemblies. For the Green's function outside the spherical assembly, we have used a simpler formula:

$$g_i^*(\boldsymbol{r}) = \frac{1}{2\pi(D_o + D_i)|\boldsymbol{r} - \boldsymbol{r_i^*}|} - \frac{(D_o - D_i)}{4\pi(D_o + D_i)^2} \int_0^1 \frac{1}{|\boldsymbol{r} - x\boldsymbol{r_i^*}|} x^{-\frac{D_i}{D_o + D_i}} dx, \tag{3}$$

To analytically calculate the Green's function values for certain $\boldsymbol{r}$ and $\boldsymbol{r_i^*}$, we used the *scipy.integrate.quad* function to compute the intergrals in Eq. 15 and Eq. S3 via python.

To evaluate the accurancy of numerical results, we have compared the Green's function values calculated via COMSOL software with the analytical results for spherical aggregates without potential of mean force, as shown in Fig. S1 B/C.

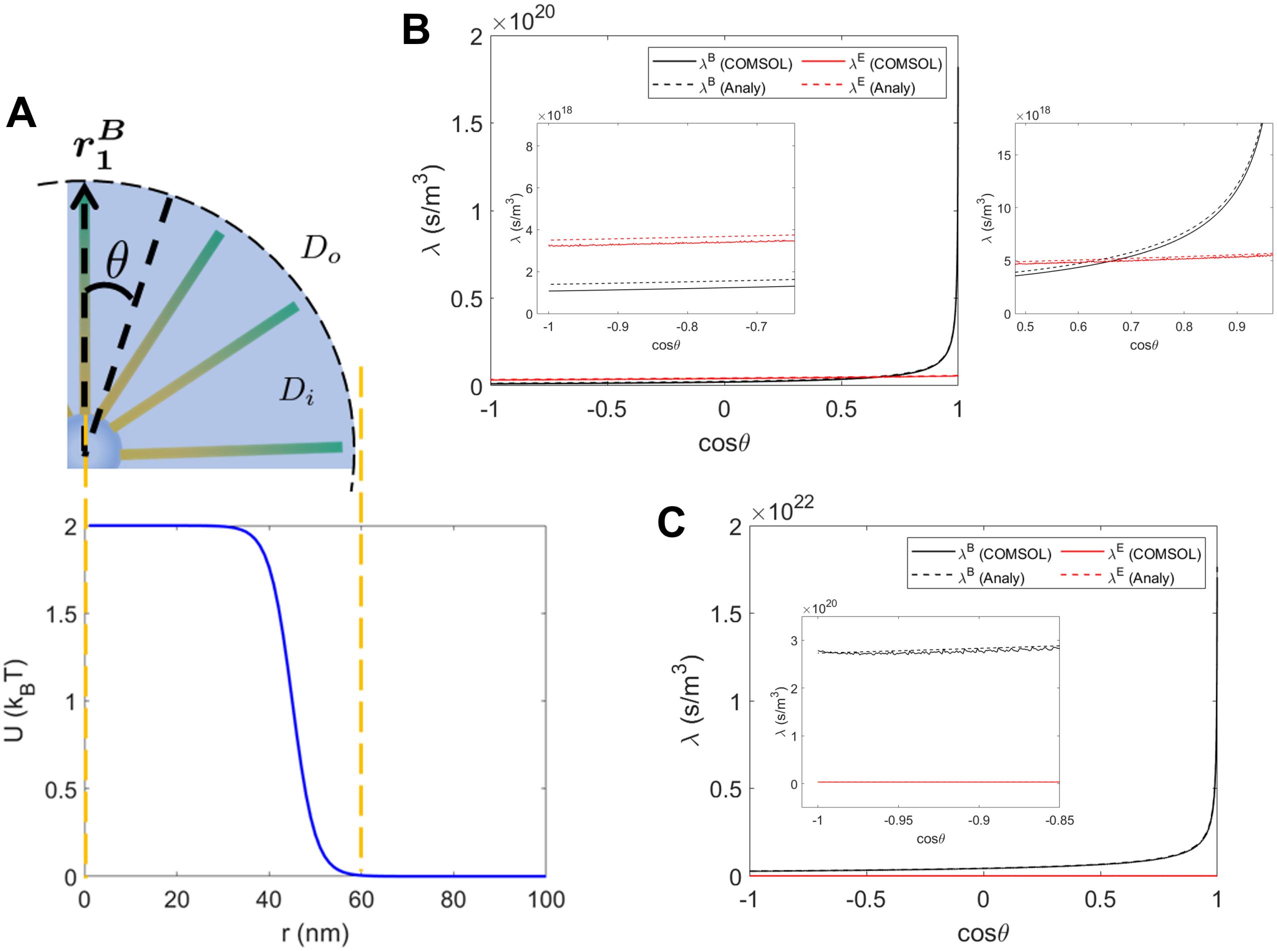


**Fig. S1.** **A**: A partial diagram showing the structure of the spherical aggregates and the function $U(r)$ with $U_0 = 2k_BT$, $\eta = 5nm$, and $r_0 = 45nm$ as the potential of mean force. $\boldsymbol{r_1^B}$ represents the vector from the center of the aggregate to the start codon of one mRNA and $\theta$ means the angle between this vector and other mRNA vectors. **B**: Green's function values at $\boldsymbol{r_1^B}$ from point sources at the start codons ($\lambda^B$, shown in black lines) or at the stop codons ($\lambda^E$, shown in red lines) of other mRNAs in spherical CPEB aggregate when $D_o/D_i = 30$. Numerical results are shown in solid lines while the analytical results are shown in dashed lines. Two regions are zoomed in to show the differences in detail. **C**: Green's function values at $\boldsymbol{r_1^B}$ from point sources at start codons ($\lambda^B$) or stop codons ($\lambda^E$) of other mRNAs in spherical Rim4 aggregate when $D_o/D_i = 30$. One region is zoomed in.

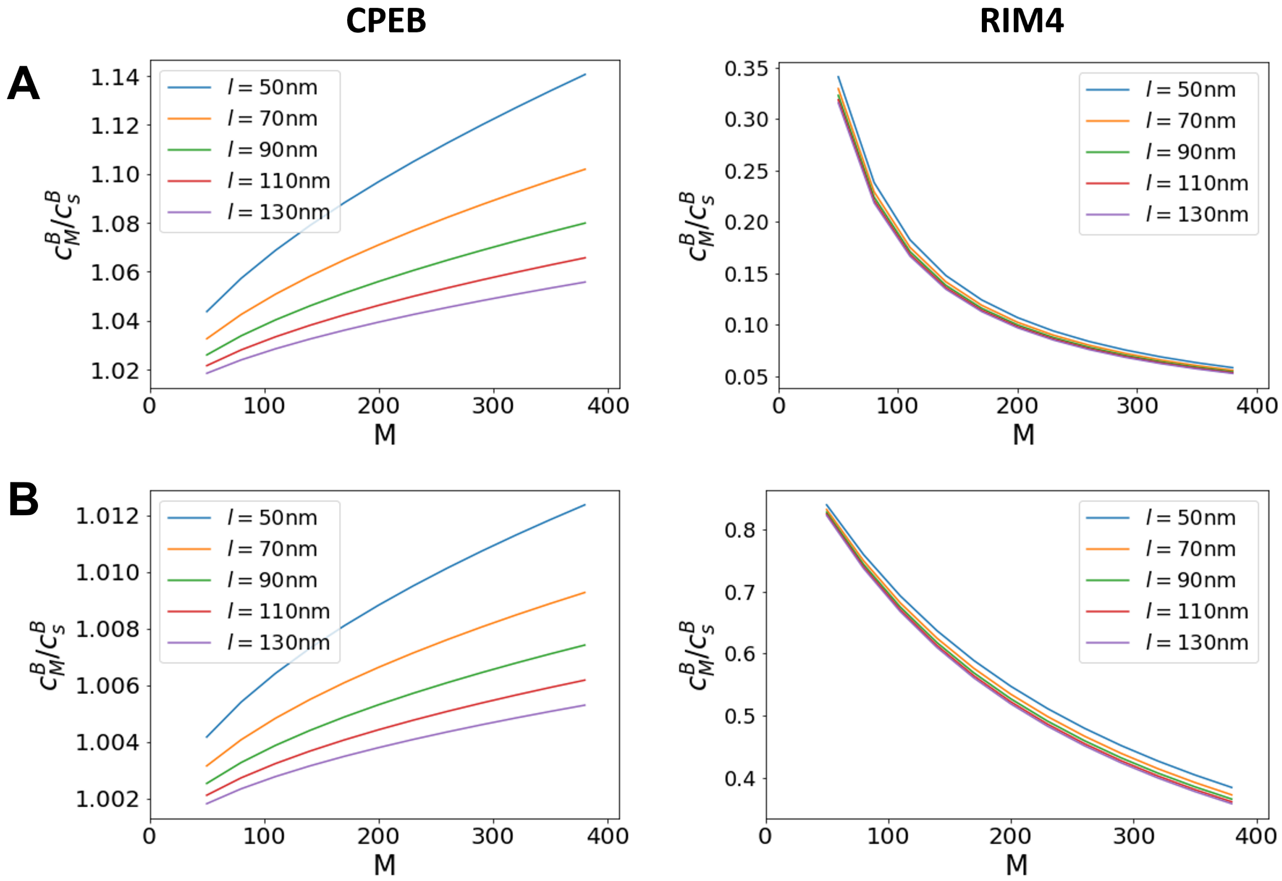


**Fig. S2.** The net translation rate for mRNAs having different lengths, $l$, in spherical aggregates with homogeneous diffusivity and no potential of mean force. **A**: Results for diffusion-limited initiation ($k_{in} = 2 \times 10^6 \ \ nm^3/s$). **B**: Results for slow-initiation ($k_{in} = 2 \times 10^5 \ \ nm^3/s$).

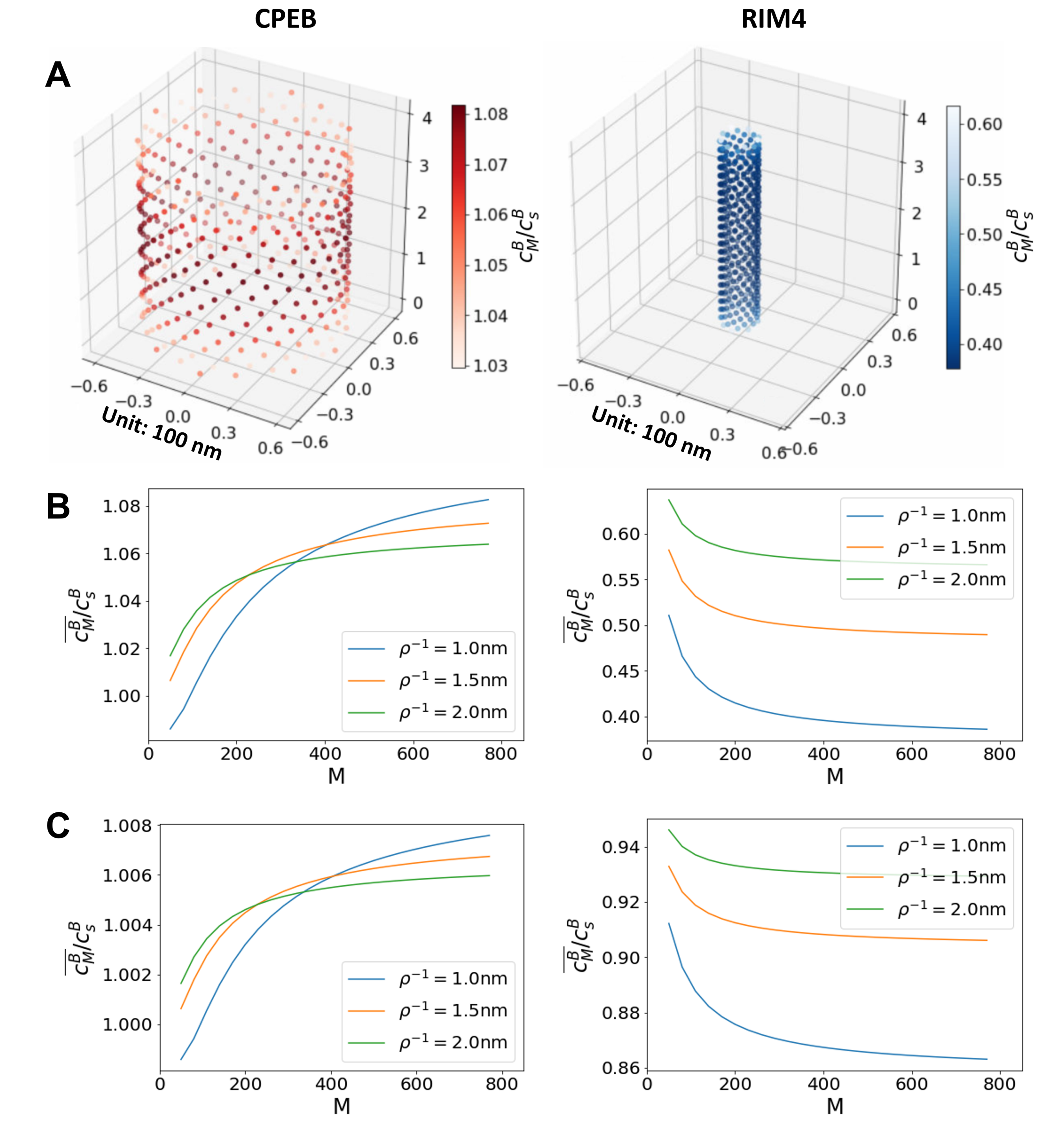


**Fig. S3.** The net translation rate in cylindrical aggregates having homogeneous diffusivity and no potential of mean force. **A**: The translation rate distribution for 400 mRNAs with a linear density $\rho$ of $1\ \ nm^{-1}$ and an initiation rate $k_{in}$ of $2 \times 10^6\ \ nm^3/s$. Here we only shown the start codons using scatter points. Translational control is more significant nearer the middle of the cylinder than at the two ends. **B**: Average translation rate is plotted against aggregate size $M$ with diffusion-limited initiation ($k_{in} = 2 \times 10^6\ \ nm^3/s$). **C**: Average translation rate with slow-initiation ($k_{in} = 2 \times 10^5\ \ nm^3/s$). Several linear density values have been tested here. The translational control is more significant for higher linear density cases except when the end effects of the cylinder dramatically reduced the channeling effect in the aggregates with small zise and high linear density. **Left panel**: Results for mRNA/CPEB aggregate. **Right panel**: Results for mRNA/Rim4 aggregate.

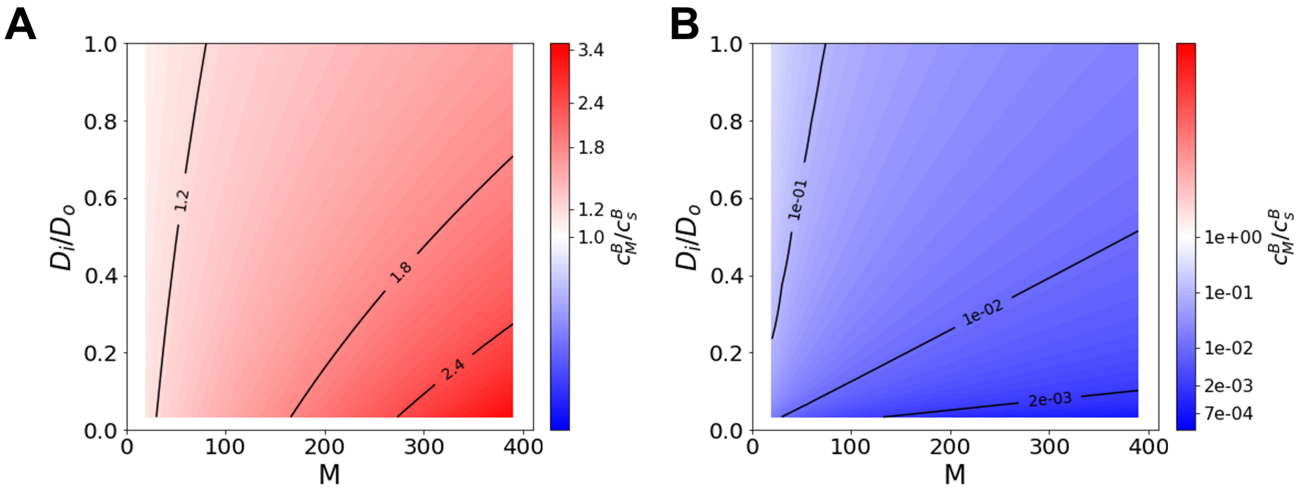


**Fig. S4.** Translation rates in spherical aggregates when $U(\boldsymbol{r}) = 0$ and $k_{in} = 6 \times 10^6 \ \ nm^3/s$ for CPEB (**A**) and for Rim4 (**B**). The color code and axes labels are the same as in Fig. 3 in the main text.

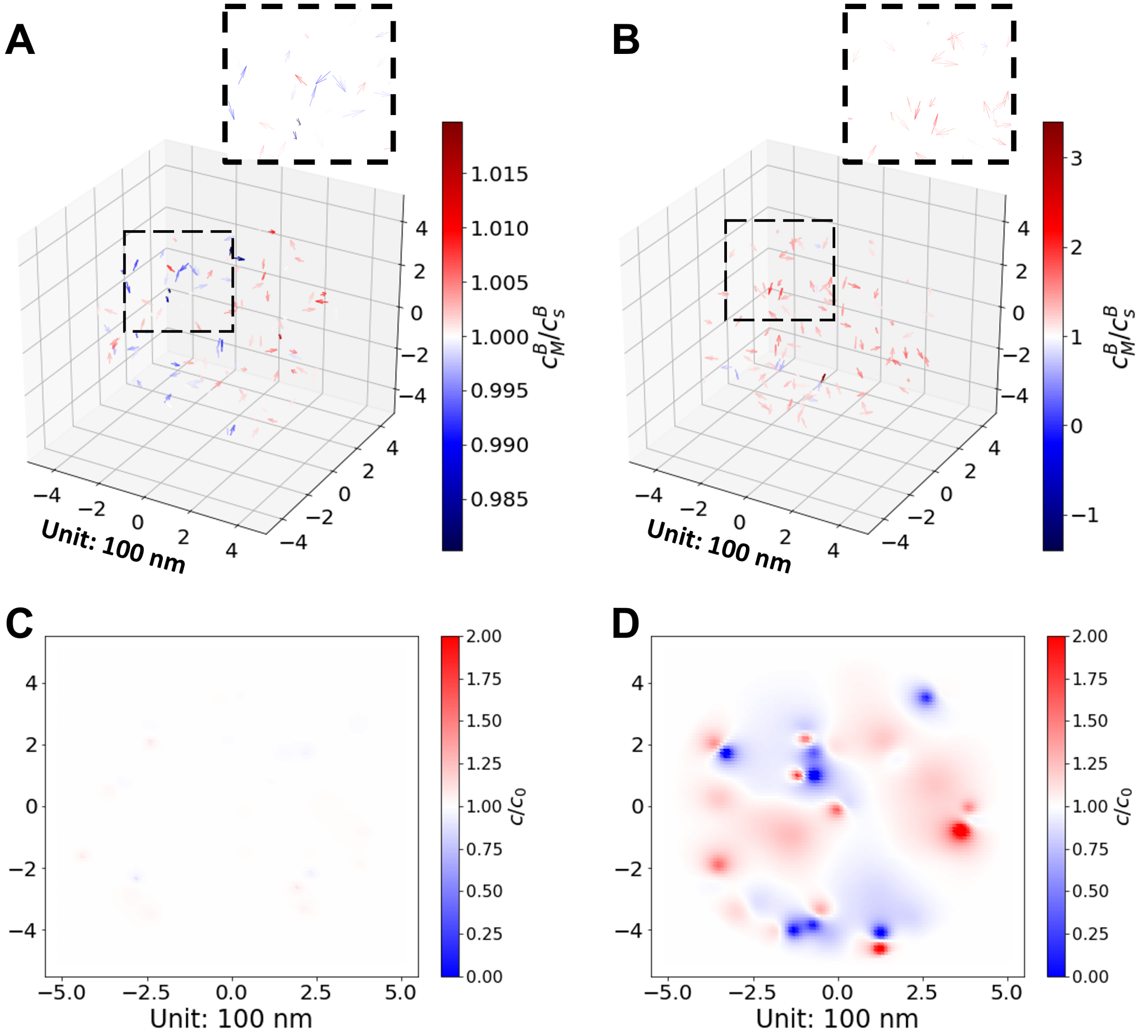


**Fig. S5.** **A, B**: The translation rate distribution coming from around 100 mRNAs in a completely random condesate with a radius of 500 nm when $U(\boldsymbol{r}) = 0$ and $k_{in} = 2 \times 10^6 \ nm^3/s$. **A**: $D_o/D_i = 1$. **B**: $D_o/D_i = 30$. Here the mRNAs are shown as vectors pointing from the start codons to the stop codons. Regions highlighted by dashed blocks are zoomed in to show detailed distributions. **C, D**: Relative ribosome concentration, $c/c_0$, at the z=0 plane in Figure **A, B**. The cutoff values are set to be 0 and 2 as minimum and maximum.

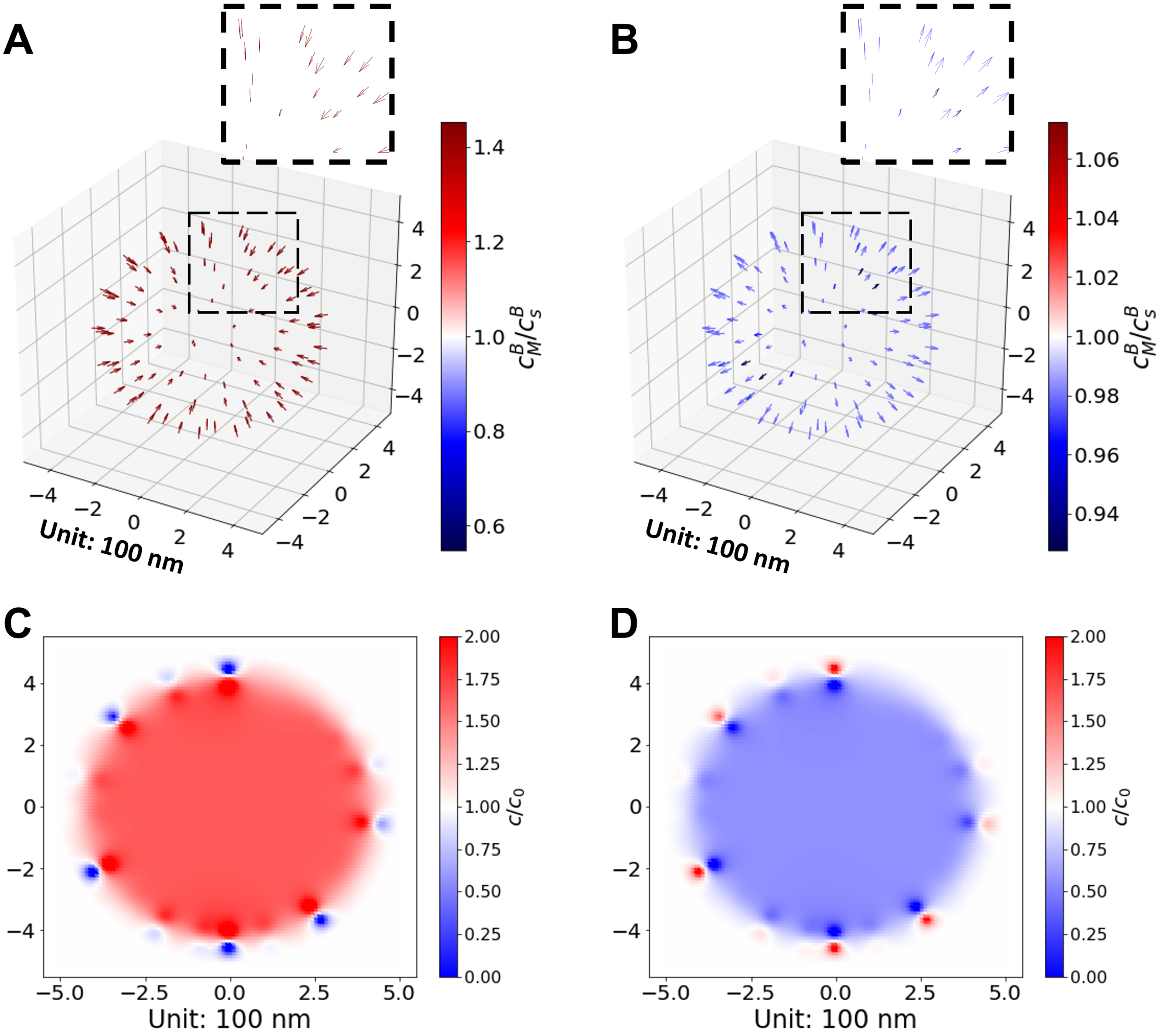


**Fig. S6.** The translation rate distribution of 100 CPEB target mRNAs in a globally polarized condesate with a radius of 500 nm when $U(\boldsymbol{r}) = 0$, $D_o/D_i = 30$ and $k_{in} = 2 \times 10^6 \ nm^3/s$. **A**: Results for a condensate with an RNA shell with an outer radius of 450 nm. **B**: Results for a condensate with an RNA core with an outer radius of 450 nm. Here the mRNAs are shown as vectors starting from the start codons to the stop codons. Regions highlighted by dashed blocks are zoomed in to show detailed distributions. **C, D**: Relative ribosome concentration, $c/c_0$, at the z=0 plane in Figure **A, B**. The cutoff values are set to be 0 and 2 as minimum and maximum.

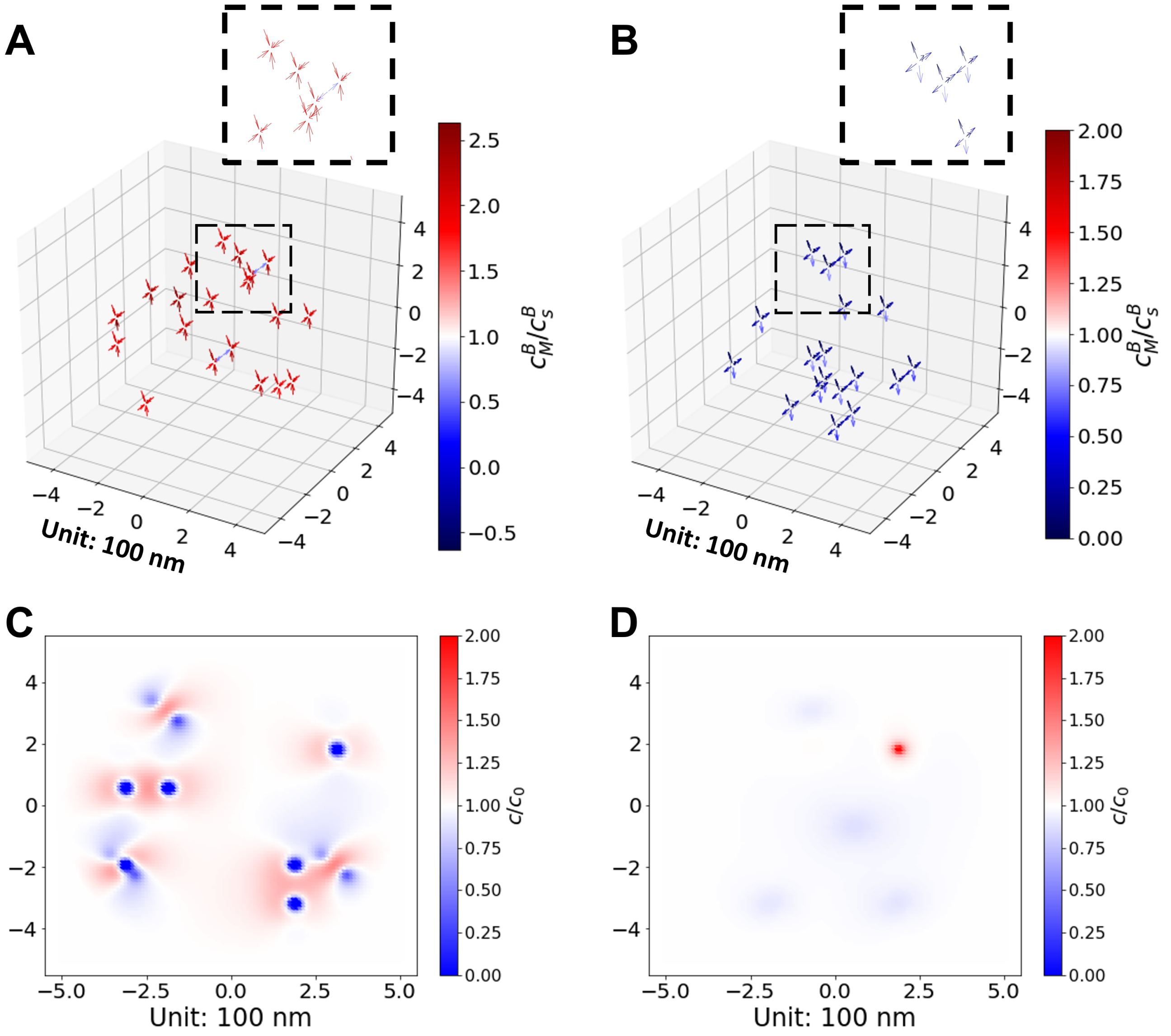


**Fig. S7.** The translation rate distribution coming from around 100 mRNAs in locally polarized condesate with a radius of 500 nm when $U(\boldsymbol{r}) = 0$, $D_o/D_i = 30$ and $k_{in} = 2 \times 10^6 \; nm^3/s$. **A**: Results for a condensate made up of randomly distributed CPEB/RNA oligomers with a size of 5. **B**: Results for a condensate made up of randomly distributed "Rim4"/RNA oligomers with a size of 5. Here the mRNAs are shown as vectors pointing from the start codons to the stop codons. Regions highlighted by dashed blocks are zoomed in to show detailed distributions. **C, D**: Relative ribosome concentration, $c/c_0$, at the z=0 plane in Figure **A, B**. The cutoff values are set to be 0 and 2 as minimum and maximum.

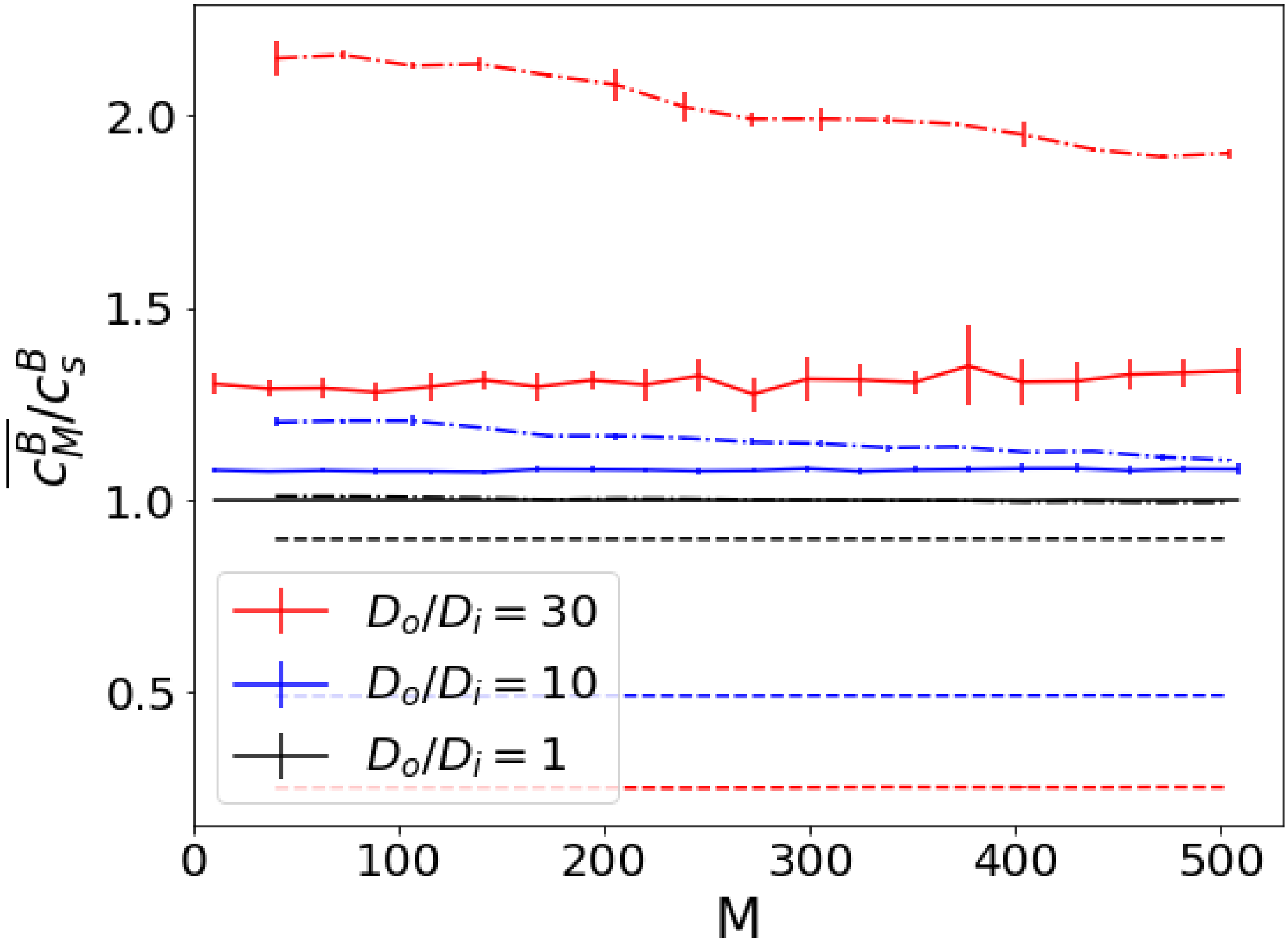


**Fig. S8.** The translation rate in locally polarized condesate when $U(\boldsymbol{r}) = 0$ and $k_{in} = 2 \times 10^6 \ \ nm^3/s$. The results for the locally polarized condensate with CPEB/RNA oligomers (shown in dot-dashed lines) or with "Rim4"/RNA oligomers (shown in dashed lines) are compared with those for the completely random condensate (shown in solide lines) when $D_o/D_i$ equals to 1 (shown as black lines), 10 (shown as blue lines) or 30 (shown as red lines). For locally polarized condesates, each oligomer has a size of 5 and five random oligomer distributions were generated for each data point to calculate the mean values and error bars.
